# Peptide ligands of the cardiac ryanodine receptor as super-resolution imaging probes

**DOI:** 10.1101/2020.06.26.131854

**Authors:** Thomas M. D. Sheard, Luke Howlett, Hannah Kirton, Zhaokang Yang, Georgina Gurrola, Derek Steele, Izzy Jayasinghe, Hector H. Valdivia, John Colyer

**Affiliations:** Faculty of Biological Sciences, University of Leeds, LS2 9JT, UK; Badrilla Ltd., Leeds Innovation Centre, Leeds LS2 9DF, UK; Cardiovascular Research Center, University of Wisconsin, Madison, WI 53705, USA; Instituto de Biotecnología, Universidad Nacional Autónoma de México, Mexico City, Mexico

**Keywords:** Peptide ligands, ryanodine receptor, RyR2, DPc10, imperacalcin, super-resolution microscopy, cardiac

## Abstract

To study the structural basis of pathological remodelling and altered calcium channel functional states in the heart, we sought to re-purpose high-affinity ligands of the cardiac calcium channel, the ryanodine receptor (RyR2), into super-resolution imaging probes. Imperacalcin (IpCa), a scorpion toxin peptide which induces channel sub-conduction states, and DPc10, a synthetic peptide corresponding to a sequence of the RyR2, which replicates arrhythmogenic CPVT functional changes, were used in fluorescent imaging experiments.

Isolated adult rat ventricular cardiomyocytes were saponin-permeabilised and incubated with each peptide. IpCa-A546 became sequestered into the mitochondria. This was prevented by treatment of the permeabilised cells with the ionophore FCCP, revealing a striated staining pattern in confocal imaging which had weak colocalisation with RyR2 clusters. Poor specificity (as an RyR2 imaging probe) was confirmed at higher resolution with expansion microscopy (proExM) (~70 nm).

DPc10-FITC labelled a striated pattern, which had moderate colocalisation with RyR2 cluster labelling in confocal and proExM. There was also widespread non-target labelling of the Z-discs, intercalated discs, and nuclei, which was unaffected by incubation times or 10 mM caffeine. The inactive peptide mut-DPc10-FITC (which causes no functional effects) displayed a similar labelling pattern.

Significant labelling of structures unrelated to RyR2 by both peptide conjugates makes their use as highly specific imaging probes of RyR2 in living isolated cardiomyocytes highly challenging.

We investigated the native DPc10 sequence within the RyR2 structure to understand the domain interactions and proposed mechanism of peptide binding. The native DPc10 sequence does not directly interact with another domain, and but is downstream of one such domain interface. The rabbit Arg2475 (equivalent to human Arg2474, mutated in CPVT) in the native sequence is the most accessible portion and most likely location for peptide disturbance, suggesting FITC placement does not impact peptide binding.

## Introduction

The ryanodine receptor (RyR), the largest known ion channel, is central to the process of excitation-contraction (EC) coupling ^1^, and has been the focus of extensive research over the last few decades. RyRs are responsible for Ca^2+^ release from the sarcoplasmic reticulum (SR) ^2^, which occurs when intracellular [Ca^2+^]i reaches a threshold concentration, in a process termed Ca^2+^-induced Ca^2+^ release (CICR) ^3^. A single homo-tetrameric, mushroom-shaped RyR consists of a cytoplasmic region which interacts with a plethora of ligands, and a transmembrane region containing the ion-conducting pore ^4^.

Advancements in microscopy have progressed the understanding that RyRs self-assemble into clusters, forming discrete functional calcium release units (also known as nanodomains). Electron microscopy gave the first glimpses of the profile of individual RyRs in peripheral couplings (interface with sarcolemmal surfaces), and in dyads (interface with t-tubule membranes) ^5–7^. More recently, electron tomography offered more detailed 3D information of cluster geometry ^8,9^.

Super-resolution techniques, which surpass the diffraction limit of light and use fluorescent markers to label targets, have revealed more insights into cluster geometry and the proteins present within these nanodomains. RyR2 clusters as imaged by confocal microscopy (limited to ~200 nm resolution) appear as smooth punctate structures, leading to over-estimations of cluster size in terms of RyRs per cluster; whereas dSTORM, which increases the resolution to ~20nm, revealed significantly smaller cluster sizes ^10–12^. The molecular-level description of peripheral nanodomains was obtained using the sub-10 nm resolution technique DNA-PAINT, sufficient to localise individual RyRs ^13^. More recently, the prevalence of RyR2 phosphorylation at the individual channel level has been studied with X10 expansion microscopy (ExM)^14^.

The efficacy of super-resolution imaging research depends on highly specific fluorescent probes, which possess high specificity for the target. Various molecules have been used as probes, such as antibodies, nanobodies, smaller protein-binding scaffolds, and peptide ligands ^15–18^. We hypothesised that two peptides with high binding specificity for the RyR, the scorpion toxin imperacalcin (IpCa) ^19^ and the synthetic DPc10 ^20^, might be effective super-resolution imaging probes, and moreover might permit the visualisation of RyRs in different functional states (open channel, IpCa; CPVT-like substate, DPc10). Reliable and highly specific non-antibody probes for this protein would also enable multiplexing, by Sheard al., 26 June 2020 – preprint copy – BioRxiv labelling other combinations of protein targets with antibodies at the same time. Furthermore, these fluorescent probes would allow understanding of possible cluster rearrangements incurred by the functional effect of each peptide.

IpCa is a 3.7 kDa peptide found in the venom of the *Pandinus imperator* scorpion ^19,21^. IpCa binds to the RyR2 with high affinity (Kd = 5-10 nM) and induces long-lasting sub-conductance states ^22,23^. The 33 amino acid peptide has an overall net charge of +8 (at physiological pH), which is clustered on one side, forming a dipole moment (Supplementary Figure 1A) ^23,24^. The exact binding site on RyR2 is unconfirmed, one study suggests IpCa enters the channel’s vestibule pore and binds an internal site ^25^, whereas a study using IpCa-biotin-streptavidin reported to bind to the cytoplasmic moiety, far from the trans-membrane pore (Supplementary Figure 1B) ^26^. Whether the peptide causes its effects by steric hindrance or an allosteric mechanism is unconfirmed ^23,27,28^. Previous imaging with a fluorescent IpCa conjugate in saponin-permeabilised cardiomyocytes confirmed an intracellular localisation in the sarcoplasm ^24^, but lacked confirmation of positive RyR2 labelling with a dual stain.

The synthetic peptide DPc10 corresponds to the RyR2 amino acid sequence Gly2460-Pro2495 (full sequence provided in methods) ^20^, and is thought to compete with the native sequence, disturbing the domain interaction. DPc10 destabilises the channel, in a way that mimics the human CPVT mutant RyR2 bearing R2474S, enhancing ryanodine binding, causing hyper-sensitisation ^20^, increasing SR Ca^2+^ leak ^29^, reducing CaM binding ^30^, without causing FKBP dissociation ^31,32^. A mutant peptide (mut-DPc10) bearing the equivalent CPVT mutation does not affect channel function ^20^, suggesting the sequence is critical for functional impact. The effect of the peptide is linked with a hypothesis made in a different peptide probe study with the peptide DP4 ^33^, which regards inter-domain interactions as being important for regulating channel function and stabilisation. This was partly informed by observations of clustered mutations in 2 domains, the N-terminal domain (1-600) and central domain (2000-2500). The native DPc10, residing in the central domain, is thought to have a partner binding region within the N-terminal domain ^34^. The DPc10 peptide disturbs this domain interaction, unzipping in order to elicit the functional effects ^20^. Previous studies using fluorescent DPc10 as an imaging probe revealed a striated labelling pattern in isolated cardiomyocytes, but lacked confirmation of RyR2 specificity with a dual-label ^35,36^.

To evaluate whether the peptides could effectively label RyR2 *in situ*, we used fluorescent conjugate forms IpCa-Alexa Fluor 546 (IpCa-A546) and DPc10-fluorescein (DPc10-FITC) to label living isolated rat ventricular cardiomyocytes, and evaluate specificity and colocalisation with confocal microscopy and protein retention ExM (proExM) ^37^. Our findings suggest that using both peptides as highly specific imaging probes for RyR2 in isolated rat cardiomyocytes is highly challenging, given poor colocalisation with RyR2 antibody labelling, and prevalence of substantial labelling of other structures in the cardiomyocyte. We also looked at high-resolution atomic models of RyR2 structure to understand the native DPc10 sequence and domain interactions, and the implications of FITC placement on peptide binding. The native sequence is within a well-structured region (helical domain 1 (HD1)). The native sequence itself is not directly interacting with an N-terminal domain, but is downstream of such a domain interface involving HD1. The native sequence is largely enclosed by neighbouring residues and inaccessible to an exogenous peptide. The rabbit Arg2475 residue (equivalent to the human Arg2474) is the most accessible part of the sequence.

## Results

### IpCa-A546 imaging

We sought to evaluate whether a fluorescent conjugate of IpCa could effectively label RyR2 in situ in isolated rat ventricular cardiomyocytes. We conjugated IpCa to Alexa Fluor 546 (Figure 1A), a fluorophore which would be compatible with the super-resolution technique proExM. The biological activity of this conjugate, in terms of retained high affinity for RyR2, was verified akin to that previously shown ^24^.

**Figure 1.**
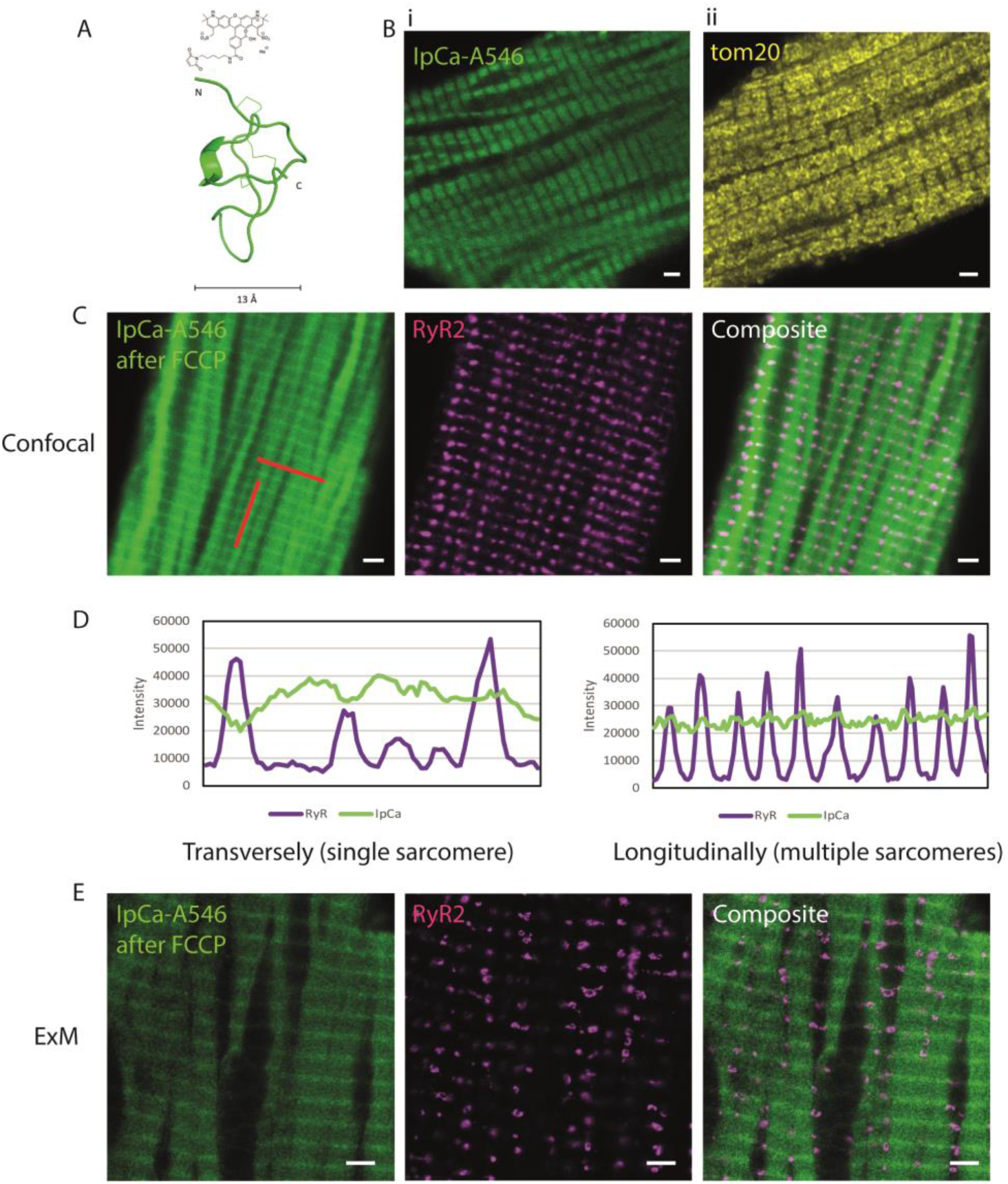
Imperacalcin-A546 as an RyR2 imaging probe in isolated rat ventricular cardiomyocytes. (A) Schematic of IpCa-A546 conjugate. (B) Isolated ventricular cardiomyocytes were incubated with 5 μM IpCa-A546 for 30 minutes. Mitochondrial sequestration was visible in confocal images (B-i), resembling the pattern typical of the mitochondrial protein tom20 (B-ii). (C) After treatment with the ionophore FCCP (30 μM for 10 minutes), IpCa-A546 uniform striated labelling was visible (green), behind the background trace of residual mitochondrial sequestration. IpCa-A546 shared a weak overlap with RyR2 labelling pattern (magenta). (D) Line profiles plots (red lines in C) revealed differences in the two labelling patterns. IpCa-A546 labelling was uniform along a single sarcomere while RyR2 varied, and across multiple sarcomeres the Z-disc peaks were less prominent in IpCa-A546 plots due to high background between Z-discs. (E) Expansion microscopy demonstrates the prevalence of non-target labelling of Z-discs, and weak colocalisation with RyR2 clusters. Scale bars, 2 μm (B,C), 8 μm (E).

Following isolation, cardiomyocytes were saponin-permeabilised to allow delivery of the peptide probe. Once attached to laminin-coated chambers, cardiomyocytes were incubated with 5 μM IpCa-A546 in a mock intracellular solution (with 10 mM caffeine and low free [Ca2+]) for 30 minutes at room temperature. Initially, IpCa-A546 was sequestered into the mitochondria, revealing a labelling pattern in confocal imaging after fixation that resembled the mitochondrial importer tom20 (Figure 1B). The reason for this sequestration is likely due to the net positive charge on the IpCa peptide ^23,24^. In order to prevent mitochondrial sequestration, permeabilised cardiomyocytes were incubated with the ionophore FCCP (30 μM, 10 mins) prior to peptide incubation. FCCP allows H^+^ ions through membranes, effectively eliminating the membrane potential of the mitochondria ^38^, which we hypothesised was the driving force for positively charged peptide accumulation in the mitochondria. By pre-treating permeabilised cardiomyocytes with FCCP, IpCa-A546 labelled in a striated pattern on top of an underlying background staining pattern throughout the cell (Figure 1C), which resembles the original mitochondrial labelling. Additionally, there was nuclear labelling (not shown).

To determine whether the striated pattern was due to RyR2 labelling, we immunolabelled the cells using RyR2 antibodies. After immunolabelling, it was evident even at ~ 250 nm resolution obtained with confocal microscopy that whilst the IpCa-A546 staining was at a similar location in the sarcomere, it stained a continuous structure visually similar to the Z-discs ^39^, unlike the punctate nature of RyR2 clusters (Figure 1C).

Line-profile plots transversely (along a single Z-disc) and longitudinally (perpendicular to the Z-discs) show the peptide labelling possesses similar Z-disc periodicity but different characteristics (Figure 1D). For instance, transversely (left), IpCa-A546 is mostly uniform, while RyR2 is variable with prominent increases in intensity at locations of clusters. Longitudinally (right), while RyR2 labelling rises and falls with clear gaps between each Z-dizc, IpCa-A546 has a Z-disc periodicity, but is much less prominent due to high (and even) cytoplasmic background in between Z-discs. These observations further suggest that IpCa-A546 striated labelling might reflect that of the Z-disc.

The degree of colocalisation between IpCa-A546 and RyR2 was assessed by thresholding 2-color confocal images and using the coloc2 Fiji plugin. Colocalisation was assessed in terms of correlation (Pearsons coefficient ranging from −1 to 1, whereby 1 is perfect correlation, 0 is no correlation, and −1 is perfect anti-correlation), and in terms of co-existence, also known as fractional overlap (Manders coefficient, ranging between 0 and 1, whereby 1 is perfect overlap, 0 is no overlap) ^40^. The Costes statistical approach states whether the result was due to chance ^41^, determining what percentage of randomised images have worse correlation than the real image. If greater than 95% (>0.95) of randomised have worse correlation, then the result is not due to chance. For instance, 1.0 or 100% means none of the randomised images had better correlation, 0.5 means 50% had better correlation, and given it is <0.95 the result could be due to chance. IpCa-A546 exhibited weak colocalisation with RyR2 in confocal data, with weak anti-correlated Pearsons coefficient −0.05 ± 0.2 suggesting the two patterns have no correlation. The fractional overlap Manders coefficient 0.08 ± 0.12 (proportion above the auto-threshold of RyR2) suggests there was almost no co-occurrence. Manders may be a less appropriate parameter given one structure (RyR2) dominates in terms of area, due to high background levels in IpCa-A546 images leading to higher thresholds and thus selecting fewer pixels. The Costes value 0.27 (<0.95) indicates these figures could be due to random chance, likely attributable to the high background/noise of IpCa images.

We used proExM to verify the degree of colocalisation at higher resolution. As suggested by confocal colocalisation analysis, most of the IpCa-A546 labelling was outside of regions of RyR2-positive labelling (Figure 1E). IpCa-A546 appeared to label the Z-discs (the continuous borders of the sarcomere), and the background traces of mitochondrial sequestration were prominent.

Thus, in our hands, IpCa-546 does not fill the criteria of a highly specific imaging probe of the RyR2 in living isolated cardiomyocytes. We also used a different fluorophore conjugate of the IpCa, with Janelia Fluor 549, but this was also heavily sequestered into mitochondria, and after FCCP showed little RyR2 colocalisation (not shown). With the present methods, IpCa appears to be ineffective as an imaging probe delivered in living cardiomyocytes, attributable largely to the significant positive charge leading to significant background binding of numerous structures in addition to the target protein RyR2.

### DPc10-FITC imaging

We next explored DPc10 as an imaging probe for RyR2. A fluorescent conjugate of DPc10 with an N-terminal 5’ carboxyfluorescein (FITC) moiety was synthesised (Figure 2A). The predicted peptide folding (using JPred software) into 2 alpha helices is like the folding of the native RyR2 sequence, as displayed for the schematic of the conjugate in Figure 2A. The FITC on the N-terminus is proposed to not inhibit peptide binding to the RyR2, given it is placed far from the Arg residue which is thought to be critical for binding, and in a solvent facing space ^20^. Placement of FITC at this site has previously been shown to preserve the peptide’s ability to bind and incur the increase to spark frequency and decrease to SR load^35^.

**Figure 2.**
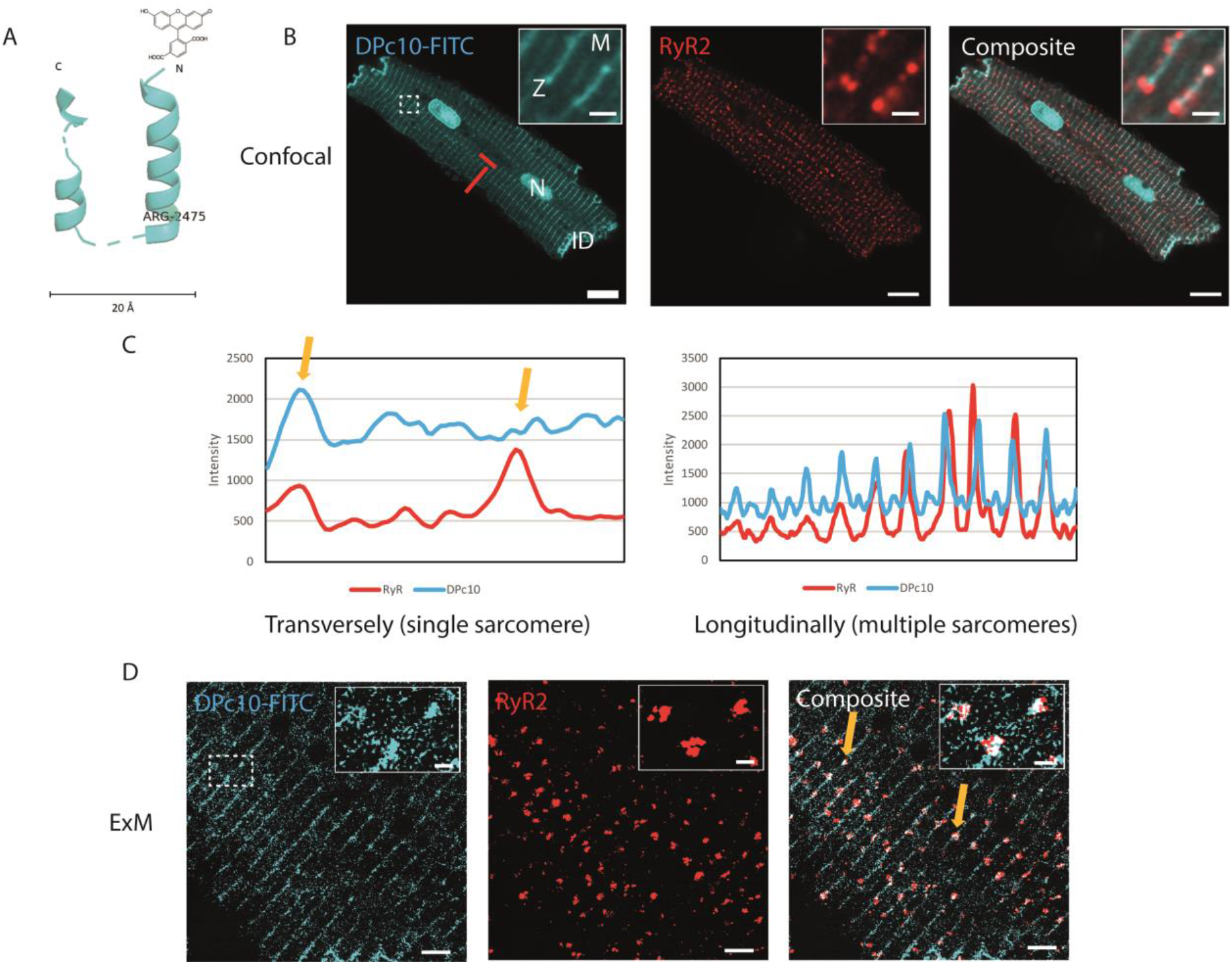
DPc10-FITC as an RyR2 imaging probe in isolated rat ventricular cardiomyocytes. (A) Schematic of DPc10-FITC conjugate, using the RyR2 structure of the native DPc10. FITC was placed at the N-terminus, far from the Arg residue critical for binding. (B) Isolated ventricular cardiomyocytes were incubated with 5 μM DPc10-FITC (cyan) for 30 minutes. The striated labelling pattern overlapped with RyR2 (red). Non-target labelling of nuclei (N), intercalated discs (ID) and M-lines (M) was visible in DPc10-FITC. (C) Line profiles plots (red lines in B) revealed differences in the two labelling patterns. Some line profile intensity changes were common between DPc10-FITC and RyR2 labelling along a single sarcomere (left arrow), although other RyR2 peaks did not appear in DPc10-FITC (right arrow). Across multiple sarcomeres the M-line labelling of DPc10-FITC is apparent. (D) Expansion microscopy suggests some positive labelling of DPc10-FITC at RyR2 clusters (arrows), as well as non-target labelling of Z-discs. Scale bars, 8 μm (B,D), 1 μm (B-inset), 2 μm (D-inset).

We incubated saponin-permeabilised isolated cardiomyocytes with 5 μM DPc10-FITC in a mock intracellular (low free [Ca2+]) solution, for 30 minutes at room temperature. After immunolabelling with antibodies for RyR2, confocal images showed that DPc10-FITC labelled in a striated pattern, which displayed moderate colocalisation with RyR2 clusters (Figure 2B). The DPc10-FITC pattern was qualitatively more punctate than IpCa-A546 (referring to the roundness of the blobs, a characteristic of RyR2 labelling). Despite the instances of colocalisation at each Z-disc, there was also a significant amount of non-RyR2 labelling of intercalated discs, nuclei, and M-lines.

Line-profile plots show peptide labelling was like RyR2 but was not identical (Figure 2C). For instance, transversely (along a single Z-disc) (left) RyR2 staining and DPc10-FITC showed similar periodicity in stain intensity (left arrow), but also instances where an increase in RyR2 intensity lacked a respective DPc10 increase (right arrow). Longitudinally (perpendicular to the Z-discs) (right), the DPc10-FITC M-line labelling is apparent as smaller peaks in between those of the Z-discs.

The colocalisation of DPc10-FITC with RyR2 was assessed, as performed previously for IpCa-A546. DPc10-FITC exhibited moderate colocalisation with RyR2 in confocal data, with a fractional overlap Manders coefficient 0.54 ± 0.08 (proportion above the auto-threshold of RyR). The labels were moderately positively correlated (Pearsons 0.45 ± 0.07), and the Costes value 1 indicates these findings are real and not due to random chance (>0.95).

Despite FITC losing a significant amount of fluorescence in the proExM protocol (and thus the resolution is poor for that channel), increased RyR2 image resolution would help assess colocalisation. proExM revealed that DPc10-FITC labelling was present at some RyR2 clusters (arrows showing white pixels in composite image), but also in larger amounts outside of RyR2 clusters, labelling Z-discs (Figure 2D).

Altogether, these results suggest that DPc10-FITC does successfully label a fraction of RyR2, but there was also a proportion of non-target labelling at sites that were not RyR2 clusters, including Z-discs, M-line, nuclei and intercalated discs.

### Alterations to improve DPc10-FITC labelling and understand non-target labelling

To explore the conditions which promote DPc10-FITC binding to RyR2, we incubated isolated cardiomyocytes with the peptide for a longer period of up to 60 minutes (Figure 3A). The longer incubation did not qualitatively affect the uniform striated labelling, and the non-target labelling remained.

**Figure 3.**
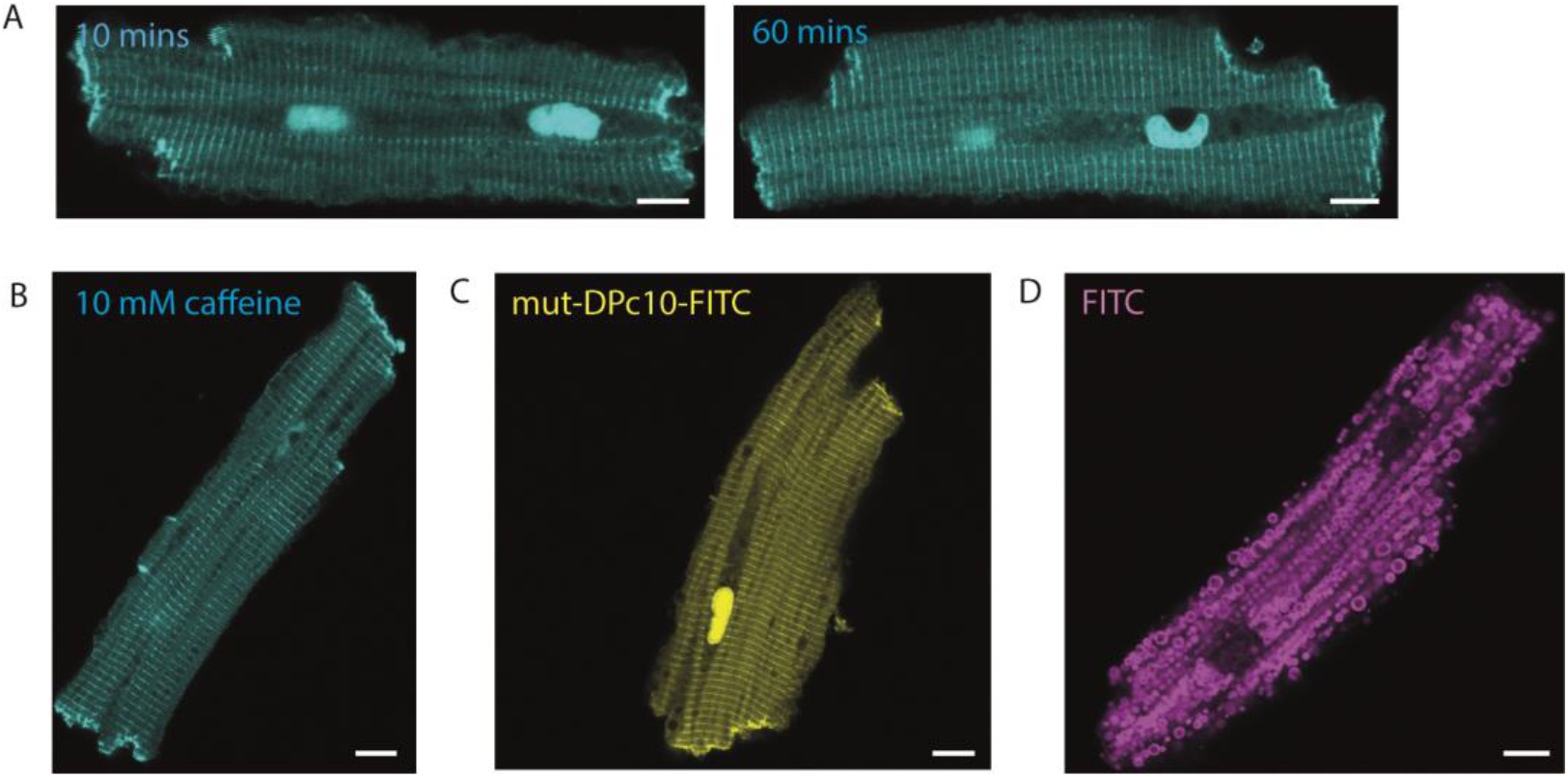
Alterations to improve DPc10-FITC target binding and understand non-target labelling. (A) Increasing the peptide incubation time to 60 minutes did not affect non-target labelling. (B) No improvement in non-target labelling was seen when including 10 mM caffeine in the peptide incubation to increase the odds of RyR2 activation. (C) 5 μM Mut-DPc10-FITC (which causes no functional alterations in RyR) produced an almost identical non-target labelling pattern. (D) 5 μM FITC (the fluorophore common to both DPc10 and mut-DPc10 conjugates) possessed an entirely different non-target labelling pattern when applied by itself, suggesting the fluorophore was not driving conjugate localisation. Scale bars, 8 μm (A-E).

We also incubated cardiomyocytes with the peptide in the presence of 10 mM caffeine (Figure 3B), which activates the channel and increases open probability ^42,43^, as has been shown to enhance access of the peptide previously ^35^. Caffeine did not qualitatively change the amount of punctate labelling, and the non-target labelling was unchanged. Whilst these results do not offer a conclusive, quantitative description of the change of % colocalisation between RyR2 and DPc10, they confirm that even with conditions optimised for RyR2 activation, there remained a significant amount of non-target labelling, which prevents DPc10-FITC being used as a specific imaging probe of RyR2 in living isolated cardiomyocytes.

To get a perspective on non-target binding, we used a similar peptide mut-DPc10, which bears the mutation equivalent to the R2475S found in CPVT, and has been shown to cause no functional defects to the RyR2 channel. The inactive peptide fails to mediate labelling of RyR2 ^34^, suggesting it loses its ability to bind RyR2. Using mut-DPc10-FITC we sought to learn whether the similar peptide-fluorophore conjugate also had non-target labelling. In confocal imaging, 5 μM mut-DPc10-FITC was like DPc10-FITC, it labelled in a striated pattern, but also had a significant amount of non-target labelling of Z-discs, nuclei and intercalated discs. (Figure 3C). This suggested that staining by these peptides does not correlate with functional data, wherein the peptides are substantially different. We explored whether the FITC fluorophore, common to both peptides, itself might influence the localisation of the peptide conjugate and account for non-target labelling. When delivered by itself, 5 μM FITC labelled an altogether different pattern, appearing to be sequestered in vesicles (Figure 3D). This suggests that the FITC fluorophore was not driving the localisation of the conjugate for subsequent staining patterns.

### RyR2 structural analysis of DPc10 native sequence, peptide binding site and domain interactions

Finally, we examined the native DPc10 sequence within various cryo-EM RyR2 structural models, PDB codes of which are detailed in the methods. We investigated whether there was a direct interaction by the native DPc10 sequence with an N-terminal domain, as suggested in the unzipping hypothesis ^20^ and alluded to by reports of peptide directly binding the N-terminal domain ^34^. We aimed to shed light on the mechanism of peptide binding and disturbing the domain interface, and to determine the implication of FITC placement on peptide binding.

Regarding the location and structure of the native DPc10 sequence (RyR2 sequence 2460-2495), it was previously described as locating in a central domain, but in modern descriptions the nomenclature has changed, and the RyR2 ‘central domain’ refers to the domain with sequence 3636-4020 ^44,45^. Therefore, DPc10 exists within helical domain region 1 (HD1), residues 2110-2679, not the ‘central domain’.

To investigate DPc10 domain interaction with the N-terminal domain and the idea that the peptide binds to the N-terminal domain directly, we looked to see whether the region containing the DPc10 sequence interacted with an N-terminal domain. The N-terminal domain of the neighbour subunit is in close proximity to HD1, but does not directly interact with the native DPc10 (~20 Å away) (Figure 4A). This N-terminal domain interacts with other residues within HD1, notably Arg2419 and Ser2426. This interaction between the N-terminal domain and HD1 is upstream of DPc10 (Figure 4B), and of the residue Arg2475 (equivalent to human Arg2474) which is mutated in CPVT. Another domain zip was observed in close proximity, involving this N-terminal domain and residues of the central domain, notably 3891-3894. We observed no intra-subunit (within the same monomer) domain interaction between the N-terminal domain and HD1 (Figure 4A).

**Figure 4.**
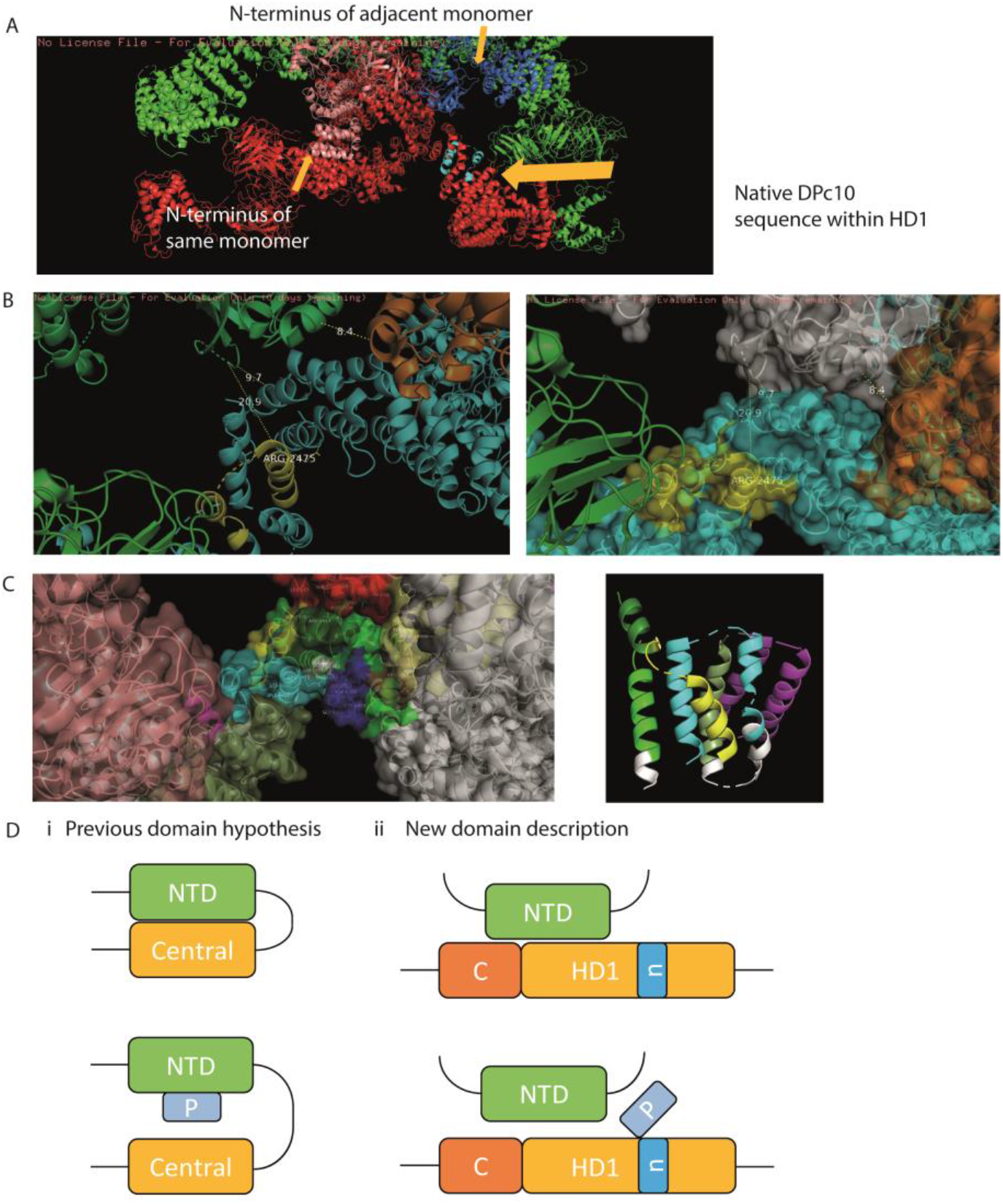
RyR2 structural analysis of the DPc10 native sequence, peptide binding site and domain interactions. (A) DPc10 native sequence exists in HD1, which comes into proximity with the neighbour subunit N-terminal domain. (B) This neighbour N-terminal domain does not interact directly with DPc10, but does interact with other residues within HD1 upstream of DPc10 and Arg2475 (in rabbit, equivalent to human Arg2474). There is also an interaction of the N-terminal domain with the central domain, in proximity to DPc10. (C) Most of the DPc10 sequence is internal and not exposed surfaces, surrounded by neighbouring helices of HD1. Arg2475 sits at the most exposed portion of the sequence. (D-i) Old schematic where peptide unzipping is by DPc10 sequence directly binding NTD. (D-ii) A proposed new schematic for the DPc10 domain zip interaction, indicating that the native DPc10 sequence, within HD1, does not directly bind to N-terminal domain, but upstream residues of HD1 do. The central domain (C) zip is also in proximity. DPc10 peptide could interact with the native sequence Arg2475, disturbing the upstream interaction of HD1 with N-terminal domain.

The simplicity of previous schematics of DPc10 peptide effect (based on direct interaction of native sequence with N-terminal domain) may be inappropriate, and we propose it may be updated to reflect that DPc10 peptide may induce domain unzipping upstream of the exact binding site, without directly binding to the N-terminal domain, instead interfering with the native sequence interactions, allosterically disturbing upstream domain zips (Figure 4D).

To gauge the likelihood and mechanism of peptide binding at the native sequence, we explored the structure and accessibility of this region. The DPc10 sequence within HD1 is largely inaccessible, surrounded by neighbouring helices, making most of the native sequence interactions internal surfaces, which were not exposed allowing disturbance by a peptide (Figure 4C). This well-structured region has a number of stabilising interactions which suggests that the degree of mobility is limited. Furthermore, this region does not become more accessible in the open confirmation, compared to the closed (Supplementary Figure 2B).

However, the native residue Arg2475 sits at the most exposed part of the native sequence and interacts with several residues within HD1, and is thus the most accessible to be disturbed by a peptide. The native Arg2475 interactions with other residues also seem to be unchanged between the open and closed conformations (Supplementary Figure 2B).

Previously, it has been shown that DPc10 peptide binding is dependent on the peptide Arg residue ^20^. A similar peptide DP4 (RyR2 sequence 2442-2477) shares one loop of the DPc10 structure, including this Arg (Supplementary Figure 2A), and similarly functionally depends on this residue ^33^, further supporting the idea that this peptide residue is critical for binding the RyR2.

To summarise on the mechanism of peptide binding near the native sequence, whilst we have no direct evidence of the site of DPc10 peptide binding, our analysis of the structure of RyR2 in the location of DPc10 sequence does not offer support to the suggestion that the peptide entirely displaces the RyR2 sequence to corrupt a domain-domain interface. We suggest the peptide may bind at the exposed DPc10 Arg2475 residue, and this is dependent on its own Arg residue. Given this concept of peptide binding, we assume that the observations made with the DPc10-FITC were not likely to have been affected by fluorophore placement at the peptide N-terminus, given this was far from the Arg residue, and would be located in solvent exposed position.

Interestingly, the DPc10 peptide binding site reported by FRET experiments is different from the location of the native DPc10 sequence within HD1, instead reporting to be around the handle region ^35^ (Supplementary Figure 2C, left). The identify the site based on a set distance 56 Å away from FKBP and CaM. In the RyR2 structure, the native sequence site is 100 Å from the location of the FKBP FRET partner (Supplementary Figure 2C, right), suggesting that this is not the site of binding they discovered, but they refer to the possibility that DPc10 may bind to the RyR2 in multiple sites.

Altogether, this information raises questions over the exact location and nature of DPc10 binding, either directly to an N-terminal domain, near the native sequence, or near the reported site based on FRET.

## Discussion

Both IpCa-A546 and DPc10-FITC possessed a significant amount of non-target labelling at sites other than the RyR2. While there were instances of positive RyR2 labelling, the prevalence of negative labelling hampers their use as high specificity super-resolution imaging probes for the RyR2 when applied to living isolated cardiomyocytes.

Despite failure to confirm specificity of the peptides for RyR2 in fluorescent imaging experiments, it is not disputed that both peptides IpCa and DPc10 bind to the RyR2. There is a wide body of evidence of the functional effects of both peptides in the literature ^20,22,24,25,29,31,34,36,46,47^.

### Evidence of positive RyR2 labelling

IpCa-A546 showed weak colocalisation with RyR2 in confocal imaging, and proExM revealed that the majority of IpCa-A546 was outside RyR2 clusters, tending to uniformly label the Z-discs which are in the same rough plane as the clusters. The colocalisation seen at confocal level was likely due to poor resolution failing to discern the close (but not identical) localisations of these two probes, attributable in part to high background of IpCa-A546 traces within the mitochondria. The pattern of IpCa-A546 near RyR2 clusters was neither punctate nor noticeably like the RyR2 pattern, instead having a uniform pattern like α-actinin of the Z-disc. Binding of IpCa, which is thought to bind the channel’s open conformation ^23^, should be encouraged when the channels activate regularly and often, as was achieved with caffeine ^42,43^.

DPc10-FITC labelling on the other hand was more punctate (a characteristic pattern of RyR2 clusters) and showed moderate colocalisation with RyR2 in confocal imaging. proExM confirmed that DPc10-FITC did successfully label some RyR2 clusters, despite a significant amount of non-target labelling of the Z-disc. Coincidence at the super-resolution level was visible, but formal quantitation was not possible due to poor resolution in the FITC channel. Future experiments with a proExM-compatible probe would be required to assess the degree of target-specific binding at the nanometer scale. Caffeine treatment did not noticeably change the uniform Z-disc-like labelling pattern, confirming that even in conditions optimized for RyR2 specificity, non-target labelling exists. It is of interest to quantify the change in RyR2 specificity by caffeine treatment in the future with a proExM-compatible conjugate, to determine whether the probe can reflect changes in RyR2 activation state.

DPc10 binding conditions are difficult to speculate on given the DPc10 binding site is not confirmed. Previously DPc10-FITC access was enhanced by increased RyR2 activation and pre-bound non-fluorescent DPc10, although this confocal analysis demonstrated striated labelling which was not verified as specific RyR2 labelling ^35^.

Given DPc10 effects do not easily reverse without long washout periods ^29,35^, peptide dissociation is unlikely to account for the amount bound to RyR2. The inclusion of caffeine should ensure the channels remain activated so peptides can re-bind even if they dissociated.

Use of a proExM-compatible mut-DPc10 conjugate, for colocalisation analysis in comparison to the active peptide, is desirable to explore whether the peptide labelling patterns correlate with functional binding data.

### Non-target labelling

It might be possible to use IpCa and DPc10 as imaging probes if the challenges of non-target labelling are circumvented. Typical immunolabelling experiments with IgG primary antibodies circumvent non-target labelling in fixed cells by performing a blocking step to eliminate sites which randomly bind epitopes. It remains to be seen whether it is possible to perform blocking to prevent sticking to these widespread structures, especially as blocking in the living cell is not possible without altering physiology and/or structure.

The positive charge of IpCa lead to significant non-target labelling due to mitochondrial sequestration. The distribution of positive charges on the peptide are critical for binding to RyR1 ^23,25,48^, so one would speculate that removing the charge to prevent this sequestration would also prevent the binding to the target RyR2.

While significant non-target labelling of IpCa-A546 and DPc10-FITC renders them ineffective as super-resolution RyR2 imaging probes in isolated cardiomyocytes, they might still prove useful in SR lipid bilayers, or cell types lacking dense contractile machinery or prevalent membrane networks, for instance atrial cardiomyocytes, or HL-1 (atrial tumour) cell line.

Previous exploits with fluorescent IpCa showed an intracellular localisation ^24^, and imaging with DPc10-FITC inferred successful RyR2 labelling by the striated labelling pattern ^35^. These results demonstrate the need to validate probe target specificity at higher resolution, and alongside an independent marker for the target of interest.

### DPc10 native sequence, peptide binding site and unzipping hypothesis

We expected to find the native DPc10 sequence directly adjacent to and interacting with an N-terminal domain, as suggested by the domain zip hypothesis ^33^. We found that the native sequence was not itself interacting with an N-terminal domain, but that a region upstream of DPc10 did interact. It is possible that this is the inter-domain zip that is important for regulating channel gating, and which is compromised in the CPVT mutated RyR2 bearing R2475S.

The fact that DPc10 did not directly interact with N-terminal domain was surprising, given the finding that DPc10 directly binds to the N-terminal domain ^34^. This study detected DPc10-labelled fluorescence in a fragment of RyR2 (after calpain digestion) which was bound by N-terminal domain antibodies. The similar peptide DP4 is also shown to bind the N-terminal domain ^49^, further supporting the notion that these peptides bound directly, as opposed to indirectly. Indirect labelling of N-terminal fragment could occur if peptides were bound to the HD1 region, which was itself bound to the N-terminal domain. Alternatively, it is possible there are multiple binding sites, both directly at the N-terminal domain, and at the native sequence location.

As well as the inter-subunit interaction with HD1, the N-terminal domain also interacted with the neighbour subunit’s central domain. It has been suggested that a region within the central domain, also known as the I domain (3722-4610), is involved in a domain interaction which is critical in regulating channel gating ^32,50^. This region, near the N-terminal domain interaction with HD1, could well be directly influenced by unzipping of these domains.

The native sequence is embedded within neighbouring helices of the HD1 region, with most of residue interactions being integral to the region, in a stable, undynamic way, which did not move drastically upon channel opening. One possibility is that the peptide (and critically the residue Arg which gives the peptide functionality) makes an interaction with a few of the residues that the native Arg does to interfere and corrupt the normal structure. The idea of this Arg being critical is supported by the complementary effects of the similar peptide DP4, which shares a similar structure, including the Arg residue critical for binding.

We highlight the discrepancy between the site of the DPc10 sequence within HD1 and the reported peptide binding site in the handle (residues 1652-2109, aka domain 3 in older nomenclature), near both FKBP and CaM ^35^. The paper does speculate that one DPc10 molecule may bind to 2 different sites, but the reason for binding at a different site to where the native sequence lies is unexplained. We are not ruling out DPc10 binding at the native sequence or at the site in the handle, away from the native sequence, given the possibility that DPc10 has multiple binding sites on the RyR2 ^35^. We suggest that the DPc10 peptide binding interaction is the focus of further research.

## Materials and Methods

### Peptides

IpCa peptide was produced as detailed previously ^51^. IpCa was modified with the cyclic imidoester 2-iminothiolane linker to introduce a sulfhydryl, which was reacted with fluorescent maleimide Alexa Fluor 546, as detailed previously ^24^.

DPc10 peptide sequence is (2460)GFCPDHKAAMVLFLDRVYGIEVQDFLLHLLEVGFLP(2495). DPc10-FITC and mut-DPc10-FITC (bearing R2474S mutation) were synthesised by Peptide Protein Research, the 5’ carboxyfluorescein located at the N-terminus of the peptide. The peptides were purified to >95% in acetonitrile and water containing 0.1% trifluoroacetic acid (TFA) by HPLC.

### Cardiomyocytes

All chemicals are sourced from Sigma-Aldrich unless specified otherwise. Animal experiments were performed according to UK Home Office approved methods. Ventricular cardiomyocytes were isolated from adult male Wistar rats using the Langendorff perfusion method as described previously ^52^. Cells were permeabilised in suspension with 10 μg/ml saponin (from Quillaja bark, S-2149), in a mock intracellular solution (concentrations in mM: EGTA, 0.5; KCl, 100; HEPES, 25; MgCl, 5.72; ATP, 5; creatine phosphate, 10. pH 7.0 with KOH) for 10 minutes at room temperature (RT) on a rocker. The free [Ca^2+^] concentration was estimated to be low (50-100 nM) given the additional EGTA buffering added to ensure contaminating Ca^2+^ was removed. Following permeabilisation, cells were centrifuged and re-suspended in intracellular solution prior to attachment to laminin-coated chambers (Thermo Fisher Scientific, 23017015). 11.7 μg/ml laminin coating was performed overnight at RT. For IpCa-A546 experiments, permeabilised cells were incubated with 30 μM FCCP (C2920) for 10 minutes prior to attachment.

Once attached, permeabilised cells were incubated with 5 μM DPc10-FITC, mut-DPc10-FITC, IpCa-A546 or FITC (diluted in intracellular solution), for 30 minutes unless stated otherwise. Where stated, 10 mM caffeine was including in the peptide incubation step. Cardiomyocytes were fixed with 2% paraformaldehyde for 10 minutes at room temperature.

### Immunolabelling

Fixed cells were permeabilised with 0.1% Triton X-100 in PBS for 10 mins and blocked with 10% normal goat serum (NGS, 10000C, Thermo Fisher Scientific) in PBS for one hour, both steps at room temperature. Primary antibodies for RyR2 (mouse monoclonal anti-RyR2 IgG1, Thermo Fisher Scientific, UK, MA3-916) were diluted to 1:200 in incubation solution ((w/v or v/v) 0.05% NaN_3_; 2% bovine serum albumin (Thermo Fisher Scientific); 2% NGS; 0.05% Triton X-100, dissolved in PBS) and incubated overnight at 4°C.

Cells were washed three times in fresh PBS at RT for 20 mins, then incubated with goat anti-mouse IgG Alexa Fluor 488 (in the case of IpCa-A546), or goat anti-mouse IgG Alexa Fluor 594 (in the case of DPc10-FITC) (A11001, A11005, Thermo Fisher Scientific).

### Imaging and expansion microscopy

For confocal imaging an inverted LSM700 (Carl Zeiss, Jena) was used with a Plan-Aprochromat 40x 1.3 NA objective. Fluorophores were excited with 488 nm and 561 nm DPSS lasers, and emission bands were selected using the in-built spectral detector.

proExM was performed as detailed in the cited paper ^37^. To image expanded gels, squares of gel were cut out and placed into acrylic chambers onto a glass no. 1.5 coverslip (Menzel Gläser), which were poly-L-lysine coated (0.1%, overnight at RT).

### Software

Image analysis was performed in Fiji (https://imagej.net/Fiji). Colocalisation analysis was implemented using the coloc2 module after performing background subtraction. A median filter was applied to DPc10-FITC ExM data to remove noise.

Pymol was used to visualize the peptides and RyR2 cryo-EM structures. The following PDB structures were used: 1ie6 (IpCa) ^48^, 5L1d (rabbit RyR2 closed) ^44^, 5goa 5go9 (porcine RyR2 open and closed) ^53^, 6ji8 (porcine RyR2 with FKBP and CaM) ^54^.

Jpred4 (http://www.compbio.dundee.ac.uk/jpred4/) was used to predict DPc10 peptide secondary structure features.

## End Matter

### Author Contributions and Notes

T.S., D.S., H.V, I.J. and J.C. designed research. H.V., G.G., H.K., Z.Y., L.H., D.S. provided materials towards primary experiments. T.S. performed experiments. T.S. and J.C. analysed data. T.S. and J.C. wrote the paper.

J.C. is a shareholder and officer of Badrilla Ltd., a company that supplies DPc10 peptides for research use only.

## Acknowledgments

The authors acknowledge the Medical Research Council funded DiMeN PhD studentship, and innovation placement funding awarded jointly to I.J. and Badrilla for covering the salary of T.S. and research costs.

We thank Badrilla for gifting DPc10-FITC, mut-DPc10-FITC, and the Valdivia lab for gifting IpCa-A546.

We thank Dr Sally Boxall and the University of Leeds Bioimaging facility for technical assistance.

---

**Supplementary Figure 1.**
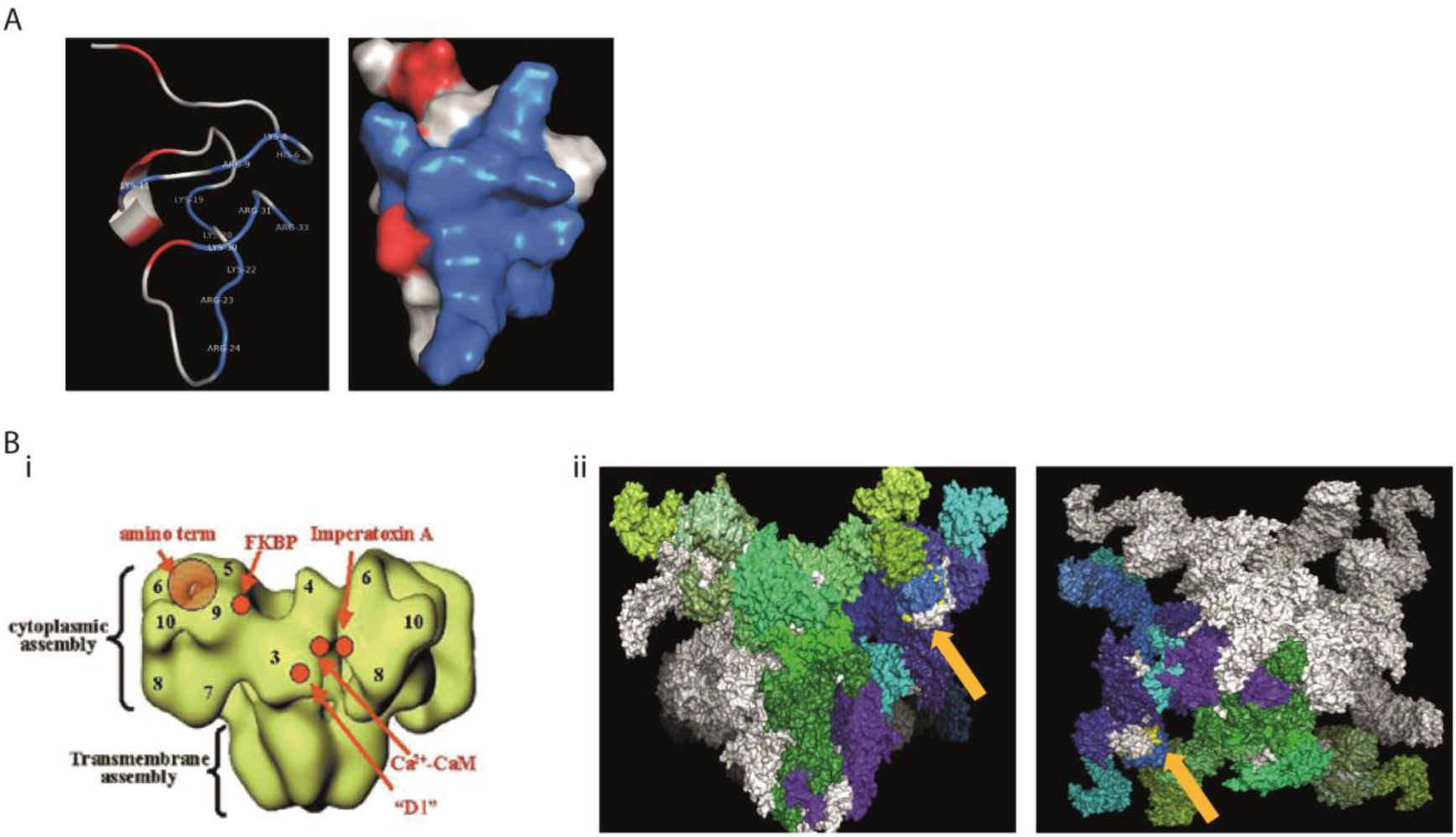
Imperacalcin background. (A) Representative image of the structure of IpCa demonstrating the cluster of positively charged residues (blue) on one side of the molecule forming the dipole moment. (Bi-ii) The exact binding site of IpCa on the RyR2 is unconfirmed, but hypothesised to be either at a cytoplasmic moiety far from the pore (annotation in B-i) or at a site within the interior of the pore (not shown).

**Supplementary Figure 2.**
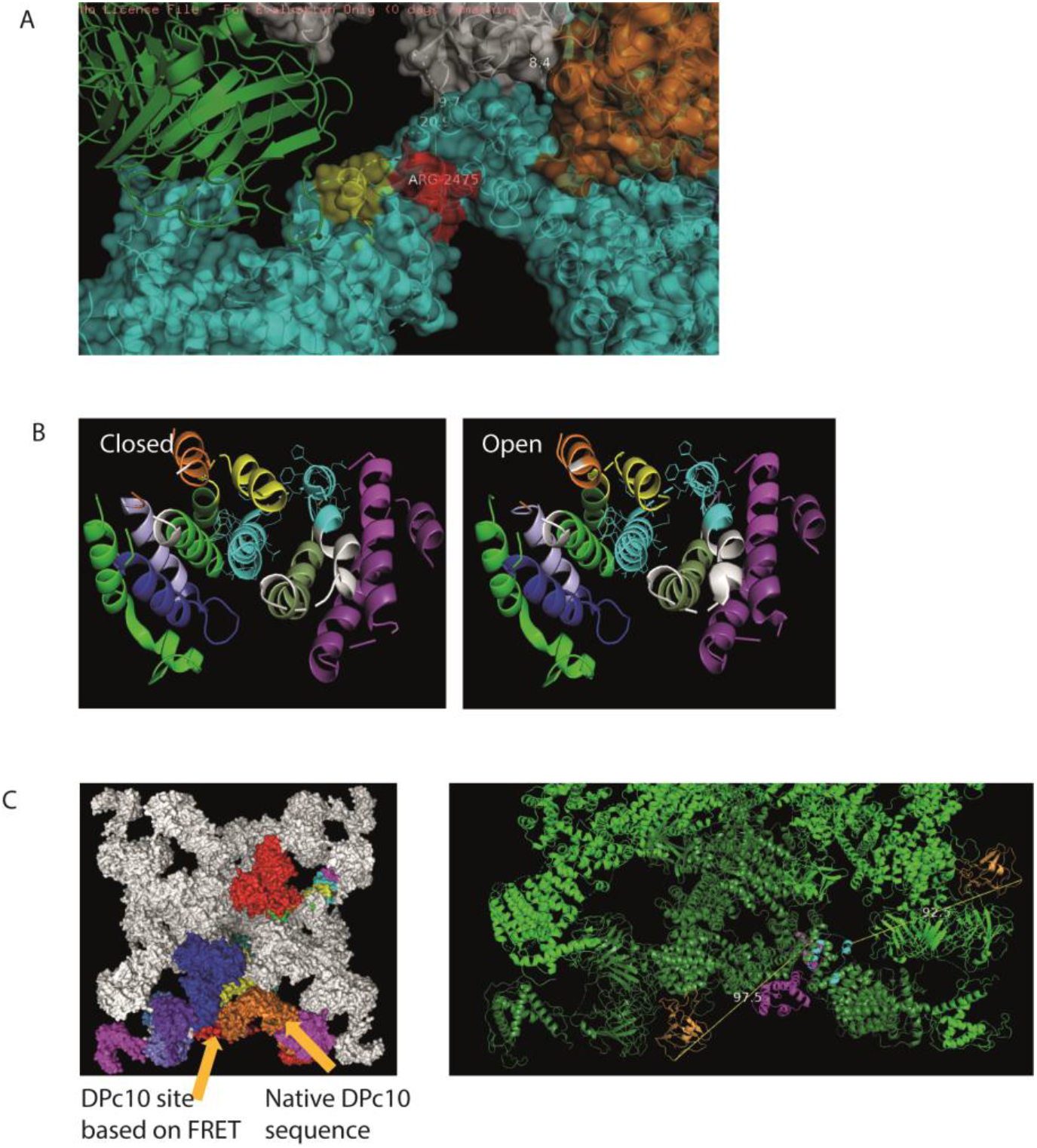
Supporting information on DPc10 sequence and peptide binding site. (A) The similar peptide DP4 relates to a similar loop/site (red) as DPc10 (yellow), which also involves Arg2475. (B) The DPc10 native sequence region becomes no more accessible in the open conformation. (C) The reported DPc10 binding site from FRET experiment (Oda 2013) is a different location to that of the native sequence region. DPc10 is reported to be distance 56 Å away from FKBP (orange) and CaM, whereas in the RyR2 structure, DPc10 native sequence is 100 Å from the location of the FKBP FRET partner.

## References

1. Stern, M. D. Theory of excitation-contraction coupling in cardiac muscle. Biophys. J. 63, 497–517 (1992).

2. Pessah, I. N., Waterhouse, A. L. & Casida, J. E. The calcium-ryanodine receptor complex of skeletal and cardiac muscle. Biochem. Biophys. Res. Commun. 128, 449–456 (1985).

3. Fabiato, A. & Fabiato, F. Contractions induced by a calcium-triggered release of calcium from the sarcoplasmic reticulum of single skinned cardiac cells. J. Physiol. 249, 469–495 (1975).

4. Takeshima, H. et al. Primary structure and expression from complementary DNA of skeletal muscle ryanodine receptor. Nature 339, 439–445 (1989).

5. Franzini-Armstrong, C. & Protasi, F. Ryanodine receptors of striated muscles: a complex channel capable of multiple interactions. Physiol. Rev. 77, 699–729 (1997).

6. Saito, A., Seiler, S., Chu, A. & Fleischer, S. Preparation and morphology of sarcoplasmic reticulum terminal cisternae from rabbit skeletal muscle. J. Cell Biol. 99, 875–885 (1984).

7. Sun, X. H. et al. Molecular architecture of membranes involved in excitation-contraction coupling of cardiac muscle. J Cell Biol 129, 659–71 (1995).

8. Asghari, P. et al. Nonuniform and variable arrangements of ryanodine receptors within mammalian ventricular couplons. Circ. Res. 115, 252–262 (2014).

9. Hayashi, T. et al. Three-dimensional electron microscopy reveals new details of membrane systems for Ca2+ signaling in the heart. J. Cell Sci. 122, 1005–1013 (2009).

10. Baddeley, D. et al. Optical single-channel resolution imaging of the ryanodine receptor distribution in rat cardiac myocytes. Proc Natl Acad Sci U A 106, 22275–80 (2009).

11. Hou, Y., Jayasinghe, I., Crossman, D. J., Baddeley, D. & Soeller, Nanoscale analysis of ryanodine receptor clusters in dyadic couplings of rat cardiac myocytes. J Mol Cell Cardiol 80, 45–55 (2015).

12. Shen, X. et al. 3D dSTORM imaging reveals novel detail of ryanodine receptor localization in rat cardiac myocytes. J Physiol (2018) doi:10.1113/jp277360.

13. Jayasinghe, I. et al. True Molecular Scale Visualization of Variable Clustering Properties of Ryanodine Receptors. Cell Rep. 22, 557–567 (2018).

14. Sheard, T. M. D. et al. Three-Dimensional and Chemical Mapping of Intracellular Signaling Nanodomains in Health and Disease with Enhanced Expansion Microscopy. ACS Nano 13, 2143–2157 (2019).

15. Harmansa, S. & Affolter, M. Protein binders and their applications in developmental biology. Dev. Camb. Engl. 145, (2018).

16. Hassanzadeh-Ghassabeh, G., Devoogdt, N., De Pauw, P., Vincke, C. & Muyldermans, S. Nanobodies and their potential applications. Nanomed. 8, 1013–1026 (2013).

17. Helma, J., Cardoso, M. C., Muyldermans, S. & Leonhardt, H. Nanobodies and recombinant binders in cell biology. J. Cell Biol. 209, 633–644 (2015).

18. Tiede, C. et al. Affimer proteins are versatile and renewable affinity reagents. eLife 6, e24903 (2017).

19. Valdivia, H. H., Kirby, M. S., Lederer, W. J. & Coronado, R. Scorpion toxins targeted against the sarcoplasmic reticulum Ca(2+)-release channel of skeletal and cardiac muscle. Proc. Natl. Acad. Sci. U. S. A. 89, 12185–12189 (1992).

20. Yamamoto, T. & Ikemoto, N. Peptide probe study of the critical regulatory domain of the cardiac ryanodine receptor. Biochem. Biophys. Res. Commun. 291, 1102–1108 (2002).

21. Zamudio, F. Z. et al. The mechanism of inhibition of ryanodine receptor channels by imperatoxin I, a heterodimeric protein from the scorpion Pandinus imperator. J. Biol. Chem. 272, 11886–11894 (1997).

22. Dulhunty, A. F., Curtis, S. M., Watson, S., Cengia, L. & Casarotto, M. G. Multiple actions of imperatoxin A on ryanodine receptors: interactions with the II-III loop ‘A’ fragment. J. Biol. Chem. 279, 11853–11862 (2004).

23. Xiao, L. et al. Structure–function relationships of peptides forming the calcin family of ryanodine receptor ligands. J. Gen. Physiol. 147, 375–394 (2016).

24. Gurrola, G. B., Capes, E. M., Zamudio, F. Z., Possani, L. D. & Valdivia, H. H. Imperatoxin A, a Cell-Penetrating Peptide from Scorpion Venom, as a Probe of Ca2+-Release Channels/Ryanodine Receptors. Pharmaceuticals 3, 1093–1107 (2010).

25. Tripathy, A., Resch, W., Le Xu, Valdivia, H. H. & Meissner, G. Imperatoxin A Induces Subconductance States in Ca2+ Release Channels (Ryanodine Receptors) of Cardiac and Skeletal Muscle. J. Gen. Physiol. 111, 679–690 (1998).

26. Samsó, M., Trujillo, R., Gurrola, G. B., Valdivia, H. H. & Wagenknecht, T. Three-Dimensional Location of the Imperatoxin a Binding Site on the Ryanodine Receptor. J. Cell Biol. 146, 493–500 (1999).

27. Ramos-Franco, J. & Fill, M. Approaching ryanodine receptor therapeutics from the calcin angle. J. Gen. Physiol. 147, 369–373 (2016).

28. Vargas-Jaimes, L. et al. Recombinant expression of Intrepicalcin from the scorpion Vaejovis intrepidus and its effect on skeletal ryanodine receptors. Biochim. Biophys. Acta 1861, 936–946 (2017).

29. Yang, Z., Ikemoto, N., Lamb, G. D. & Steele, D. S. The RyR2 central domain peptide DPc10 lowers the threshold for spontaneous Ca2+ release in permeabilized cardiomyocytes. Cardiovasc. Res. 70, 475–485 (2006).

30. Uchinoumi, H. et al. CaMKII-dependent phosphorylation of RyR2 promotes targetable pathological RyR2 conformational shift. J. Mol. Cell. Cardiol. 98, 62–72 (2016).

31. Laver, D. R., Honen, B. N., Lamb, G. D. & Ikemoto, N. A domain peptide of the cardiac ryanodine receptor regulates channel sensitivity to luminal Ca2+ via cytoplasmic Ca2+ sites. Eur. Biophys. J. EBJ 37, 455–467 (2008).

32. Tateishi, H. et al. Defective domain-domain interactions within the ryanodine receptor as a critical cause of diastolic Ca2+ leak in failing hearts. Cardiovasc. Res. 81, 536–545 (2009).

33. Yamamoto, T., El-Hayek, R. & Ikemoto, N. Postulated Role of Interdomain Interaction within the Ryanodine Receptor in Ca2+ Channel Regulation. J. Biol. Chem. 275, 11618–11625 (2000).

34. Oda Tetsuro et al. Defective Regulation of Interdomain Interactions Within the Ryanodine Receptor Plays a Key Role in the Pathogenesis of Heart Failure. Circulation 111, 3400–3410 (2005).

35. Oda, T. et al. Binding of RyR2 “Unzipping” Peptide in Cardiomyocytes Activates RyR2 and Reciprocally Inhibits Calmodulin Binding. Circ. Res. 112, 487–497 (2013).

36. Uchinoumi, H. et al. Catecholaminergic polymorphic ventricular tachycardia is caused by mutation-linked defective conformational regulation of the ryanodine receptor. Circ. Res. 106, 1413–1424 (2010).

37. Tillberg, P. W. et al. Protein-retention expansion microscopy of cells and tissues labeled using standard fluorescent proteins and antibodies. Nat Biotechnol 34, 987–92 (2016).

38. Dedkova, E. N. & Blatter, L. A. Measuring mitochondrial function in intact cardiac myocytes. J. Mol. Cell. Cardiol. 52, 48–61 (2012).

39. Jayasinghe, I., Crossman, D. J., Soeller, C. & Cannell, M. B. A new twist in cardiac muscle: dislocated and helicoid arrangements of myofibrillar z-disks in mammalian ventricular myocytes. J. Mol. Cell. Cardiol. 48, 964–971 (2010).

40. Manders, E. M.M., Verbeek, F. J. & Aten, J. A. Measurement of co-localization of objects in dual-colour confocal images. J. Microsc. 169, 375–382 (1993).

41. Costes, S. V. et al. Automatic and quantitative measurement of protein-protein colocalization in live cells. Biophys. J. 86, 3993–4003 (2004).

42. Rousseau, E., Ladine, J., Liu, Q.-Y. & Meissner, G. Activation of the Ca2+ release channel of skeletal muscle sarcoplasmic reticulum by caffeine and related compounds. Arch. Biochem. Biophys. 267, 75–86 (1988).

43. Smith, G. L. & Steele, D. S. Measurement of SR Ca2+ content in the presence of caffeine in permeabilised rat cardiac trabeculae. Pflüg. Arch. 437, 139–148 (1998).

44. Dhindwal, S. et al. A cryo-EM-based model of phosphorylation- and FKBP12.6-mediated allosterism of the cardiac ryanodine receptor. Sci. Signal. 10, (2017).

45. Samsó, M. A guide to the 3D structure of the ryanodine receptor type 1 by cryoEM. Protein Sci. Publ. Protein Soc. 26, 52–68 (2017).

46. Porta, M., Diaz-Sylvester, P. L., Nani, A., Ramos-Franco, J. & Copello, J. A. Ryanoids and imperatoxin affect the modulation of cardiac ryanodine receptors by dihydropyridine receptor Peptide A. Biochim. Biophys. Acta BBA - Biomembr. 1778, 2469–2479 (2008).

47. Terentyev Dmitry, Viatchenko-Karpinski Serge, Valdivia Héctor H., Escobar Ariel L. & Györke Sandor. Luminal Ca2+ Controls Termination and Refractory Behavior of Ca2+-Induced Ca2+ Release in Cardiac Myocytes. Circ. Res. 91, 414–420 (2002).

48. Lee, C. W. et al. Molecular basis of the high-affinity activation of type 1 ryanodine receptors by imperatoxin A. Biochem. J. 377, 385–394 (2004).

49. Kobayashi, S. et al. Dantrolene stabilizes domain interactions within the ryanodine receptor. J. Biol. Chem. 280, 6580–6587 (2005).

50. George, C. H. et al. Ryanodine receptor regulation by intramolecular interaction between cytoplasmic and transmembrane domains. Mol. Biol. Cell 15, 2627–2638 (2004).

51. Zamudio, F. Z. et al. Primary structure and synthesis of ImperatoxinA (IpTxa), a peptide activator of Ca2+ release channels/ryanodine receptors. FEBS Lett. 405, 385–389 (1997).

52. Yang, Z. & Steele, D. S. Effects of cytosolic ATP on spontaneous and triggered Ca(2+)-induced Ca(2+) release in permeabilised rat ventricular myocytes. J. Physiol. 523, 29–44 (2000).

53. Peng, W. et al. Structural basis for the gating mechanism of the type 2 ryanodine receptor RyR2. Science 354, (2016).

54. Gong, D. et al. Modulation of cardiac ryanodine receptor 2 by calmodulin. Nature (2019) doi:10.1038/s41586-019-1377-y.

